# Internal amino acid state modulates yeast taste neurons to support protein homeostasis in *Drosophila*

**DOI:** 10.1101/187310

**Authors:** Kathrin Steck, Samuel J. Walker, Pavel M. Itskov, Célia Baltazar, Carlos Ribeiro

## Abstract

To optimize fitness, animals must dynamically match food choices to their current needs. For drosophilids, yeast fulfils most dietary protein and micronutrient requirements. While several yeast metabolites activate known gustatory receptor neurons (GRNs) in *Drosophila melanogaster*, the chemosensory channels mediating yeast feeding remain unknown. Here we identify a class of proboscis GRNs required for yeast intake, and show that these GRNs act redundantly to mediate yeast feeding. While nutritional and reproductive states synergistically increase yeast appetite, we find a separation of these state signals at the level of GRN responses to yeast: amino acid but not mating state enhances yeast GRN gain. The sensitivity of sweet GRNs to sugar is not increased by protein deprivation, providing a potential basis for protein-specific appetite. The emerging picture is that different internal states act at distinct levels of a dedicated gustatory circuit to elicit nutrient-specific appetites towards a complex, ecologically relevant protein source.

## INTRODUCTION

Decision-making is a key function of the brain. One of the most ancestral and consequential decisions animals need to make is which foods to eat, since balancing the intake of multiple classes of nutrients is critical to optimizing lifespan and reproduction^1^. To do this, many animals, including humans, develop so-called “specific appetites”, seeking out and consuming specific foods in response to a physiological deficit of a particular nutrient^2–5^. Recently, several populations of central neurons driving consumption of specific nutrients have been identified in different species^6–9^. How these circuits modulate sensory processing to elicit state-specific behavioral responses, however, is poorly understood. The ability to precisely control the intake of dietary proteins is emerging as a conserved phenomenon across phyla. Insects, for example, tightly regulate their intake of protein depending on their internal states^10,11^. Mosquito disease vectors impose a huge burden on human health due to their need for dietary protein, which drives host-seeking and feeding behaviors only during specific internal states^12,13^. Dietary protein homeostasis is not specific to invertebrates, as humans are also able to select high-protein foods when low on protein^14,15^. Although protein is essential for sustaining key physiological processes such as reproduction, excessive protein intake has detrimental effects on aging and health^16–20^. This emphasizes the importance of this tight control of protein intake.

Most *Drosophila* species, including the model organism *Drosophila melanogaster*, are highly adapted to consume yeast as the major source of non-caloric nutrients in the wild, including proteins, and thus amino acids (AAs)^21–23^, as well as sterols, vitamins etc.^24^. It is therefore essential for flies to precisely regulate the intake of yeast. This is achieved by modulating decision-making at different scales, from exploration to feeding microstructure, according to different internal state signals, including AA state, reproductive state, and commensal bacteria composition^25,26^. Deprivation from dietary protein or essential AAs leads to a compensatory yeast appetite, thought to be mediated by direct neuronal nutrient sensing^20,26–28^. Activation of small cluster of dopaminergic neurons by AA deficit is involved in stimulating this appetite^9^; while in parallel, a protein-specific satiety hormone, FIT, is secreted by the fat body in the fed state and inhibits intake of protein-rich food^8^. Mating further enhances the yeast appetite of protein-deprived females through the action of Sex Peptide on a dedicated neuronal circuit^29,30^. Finally, specific commensal bacteria can suppress the yeast appetite generated by deprivation from essential amino acids, through a mechanism likely to be distinct from simply providing the missing amino acids^26^. How the nervous system integrates these different internal state signals, and how these signals modulate neural circuits that process chemosensory information and control feeding behavior, is unknown.

Contact chemosensation is critical for assessing the value of food sources for feeding^31,32^, as well as for a range of other behaviors^33–40^. As in many insects, *Drosophila* gustatory receptor neurons (GRNs) are distributed in different parts of the body^41^, and their central projections are segregated according to the organ of origin^42^. Furthermore, different classes of GRNs are thought to be specialized to detect different categories of taste stimuli, including bitter compounds, sugars, water and sodium^42–51^. Of these, the most well-characterized are sweet-sensing GRNs, which mediate attractive responses to sugars^42–44,48^, fatty acids^52,53^ and glycerol^54^. Recent work has demonstrated that sweet GRNs innervating labellar taste sensilla and pharyngeal taste organs act in parallel to support sugar feeding, since loss of either alone does not abolish flies’ feeding preference for sugar, but silencing of both drastically reduces sugar preference^55^. Specific roles have been demonstrated for these distinct sweet-sensing GRN populations: sensillar GRNs are important for initiation of feeding, suppression of locomotion^38,39^, and inhibition of egg-laying^33,36^, while pharyngeal GRNs are important for sustaining sugar feeding^55,56^ and inducing local search behavior^57^. These different classes of neurons project to discrete regions within the subesophageal zone (SEZ) in the brain of the adult fly^42^, providing a potential substrate for encoding different gustatory categories. However, the involvement of the gustatory system in yeast feeding is currently unknown.

In order to dissect the neural circuit mechanisms by which internal states homeostatically regulate protein intake, it is crucial to identify the sensory inputs that drive intake of yeast. While multiple volatiles produced by yeast fermentation are detected by the olfactory system and are highly attractive at long ranges to fruit flies^58–60^, the sensory channels that mediate feeding on yeast are unknown. Several yeast metabolites have been shown to activate subsets of taste receptor neurons in *Drosophila*. Sensillar GRNs detect glycerol, a sugar alcohol produced by yeast, through the Gr64e receptor^54^, while some amino acids have been shown to activate a subset of *Ir76b*-expressing GRNs on the legs^61,62^. Taste peg GRNs, meanwhile, have been shown to respond to carbonation, a major byproduct of alcoholic fermentation^63^, although whether this contributes to feeding has never been tested. Whether and to what extent these individual gustatory cues contribute to the high phagostimulatory power of yeast, however, is currently unknown. Thus, a major gap in our understanding of the neuroethology of *Drosophila* is that the specific sensory inputs that drive feeding on yeast, whether external or postingestive, remain to be characterized.

In this study, we show that a subset of GRNs on the proboscis is required for yeast feeding behavior. Acute silencing of these neurons drastically reduces feeding on yeast. We show that within this population of GRNs, neurons in distinct anatomical locations show physiological responses to the taste of yeast, and that these subsets act redundantly to support yeast feeding. We further demonstrate that the response of these sensory neurons to yeast is modulated by the internal AA state of the fly: deprivation from dietary AAs increases both yeast feeding and the gain of sensory neuron responses. This effect is specific to yeast GRNs, as sweet-sensing neurons are not sensitized by yeast deprivation. Furthermore, while reproductive state modulates yeast feeding behavior, it has no effect on sensory responses, indicating that distinct internal states act at different levels of sensory processing to modulate behavior. This study therefore identifies gustatory neurons that are required for the ingestion of an ethologically and ecologically key resource of the fly and identifies a circuit mechanism that could contribute to the homeostatic regulation of protein intake.

## RESULTS

### Identification of sensory neurons underlying yeast feeding

Yeast is a key nutrient source in the ecology of *Drosophila* species^22,23^. While sugars provide flies with energy, yeast is the primary source of non-caloric nutrients for the adult fly, and particularly of amino acids (AAs) and proteins. As such, flies can independently regulate their intake of sugars and yeast depending on their internal state in order to compensate for nutritional deficiencies. Deprivation from AAs specifically increases feeding on yeast, whereas deprivation from dietary carbohydrates elicits a specific increase in feeding on sucrose (Figure 1A). Though much is known about the gustatory pathways by which flies regulate intake of sugars^42–44,48,55,56,64–66^, the sensory basis of yeast feeding is not known.

**Figure 1.**
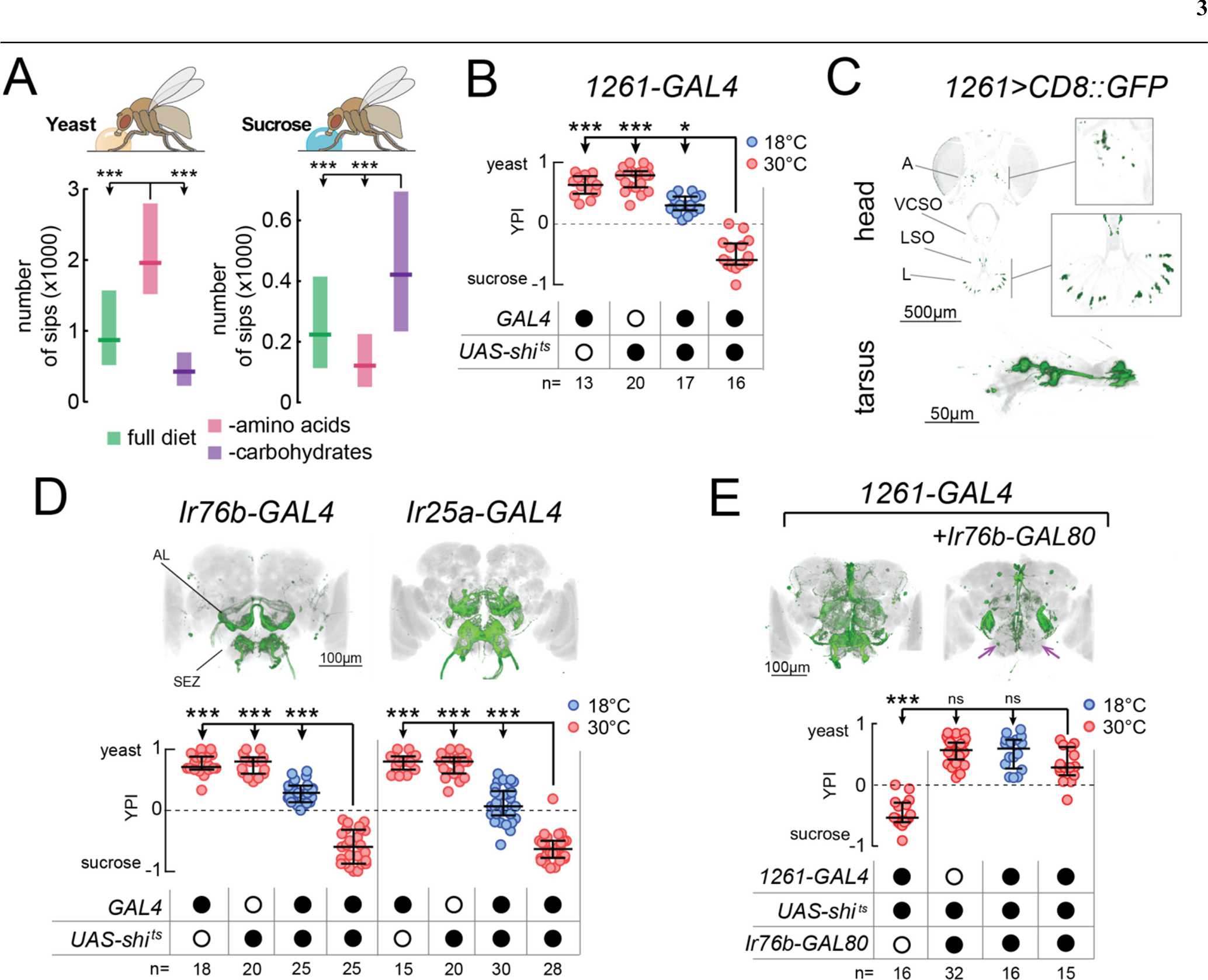
Identification of neuronal populations required for yeast feeding. (A) Number of sips from 10% yeast (left) and 20 mM sucrose (right) by mated female flies of the genotype *Ir76b-GAL4,UAS-GCaMP6s*, fed for 3 days on a holidic diet with the indicated composition. (B) Yeast preference index (YPI) of yeast-deprived female flies in which *1261-GAL4* neurons were acutely silenced and corresponding controls. (C) Expression pattern of *1261-GAL4* in the head and tarsus of the fly. Boxes are magnifications of the antenna or the labellum. Note absence of signal in the maxillary palps. Green represents GFP signal and gray the cuticular autofluorescence. (D) YPI of yeast-deprived female flies in which *Ir76b*‐ or *Ir25a-GAL4* neurons were acutely silenced and corresponding controls. (E) YPI of yeast-deprived female flies in which *1261-GAL4* neurons were all silenced, or with *Ir76b-GAL80*, and corresponding controls. Arrows indicate loss of GFP expression in the SEZ. (D) and (E) Expression pattern of experimental flies in the brain as visualized using *UAS-CD8::GFP* in green, with nc82 synaptic staining in gray. A, antenna; VCSO, ventral cibarial sense organ; LSO, labral sense organ; L, labellum; AL, antennal lobes; SEZ, subesophageal zone. In this and following figures, empty and filled black circles represent absence and presence of the indicated elements, respectively. In (A), boxes represent median with upper/lower quartiles. In (B), (D) and (E), circles represent yeast preference in single assays, with line representing the median and whiskers the interquartile range. ***p<0.001, **p<0.01, *p<0.05, ns p≥0.05. Groups compared by Kruskal-Wallis test, followed by Dunn’s multiple comparison test.

Several yeast metabolites, such as glycerol, carbonation and polyamines, have been shown to activate chemosensory neurons of *Drosophila*^37,54,62,63^. As such, we first asked whether these metabolites can account for the high phagostimulatory power of yeast in protein-deprived flies. We found that while yeast induced an extremely high level of feeding, these other substrates evoked only a very low level of feeding (Figure 1 – figure supplement 1A). We also found that deactivating yeast, such that it cannot generate carbonation, did not affect the high level of feeding on this substrate. Furthermore, silencing gustatory neurons known to respond to carbonation and glycerol did not affect yeast preference, reinforcing the idea that individually, these specific yeast metabolites do not explain the high phagostimulatory power of yeast (Figure 1 – figure supplement 1B and C). For these reasons, we devised a strategy to identify neurons required for feeding on yeast, flies’ natural food source.

We set out to isolate neurons which, when their synaptic output is acutely blocked, would abolish the high feeding preference of protein-deprived females for yeast (Figure 1 – figure supplement 2A). To this end, we employed a two-color food choice assay in which protein-deprived flies had the choice of eating sucrose or yeast^27^ to conduct an unbiased silencing screen of enhancer-trap lines (Figure 1 – figure supplement 2B and C). We identified one line (*1261-GAL4*) which showed a strong reduction in yeast preference compared to genetic and temperature controls (Figure 1B) and labelled subsets of chemosensory neurons in the head and legs (Figure 1C). As such, we designed a targeted follow-up screen in which we silenced subsets of chemosensory and neuromodulatory neurons (Supplementary Table 1) and recovered two more lines that reduced the yeast preference of protein-deprived females when their output was acutely blocked during food choice: *Ir76b-GAL4* and *Ir25a-GAL4* (Figure 1D). As with *1261-GAL4*, these lines labelled subsets of chemosensory neurons in the head and legs (Figure 1 – figure supplement 2D-F).

The loss of yeast preference upon silencing the neurons labeled in these lines could be due to a reduction in yeast feeding, and/or to an increase in sucrose feeding. To disambiguate these possibilities, we used the ability provided by the flyPAD to monitor the feeding of single flies on individual substrates with a resolution of single sips^67^. We found that in protein-deprived flies, silencing any of these *GAL4* lines specifically reduced yeast feeding, without increasing sucrose feeding, confirming that these neurons are required for feeding on yeast (Figure 1 – figure supplement 2G). To further test the specificity of this manipulation to yeast, we deprived flies from either AAs or carbohydrates, using a holidic diet, and subsequently measured feeding on the flyPAD. Silencing *1261-GAL4* reduced yeast feeding in AA-deprived flies, but did not reduce sucrose feeding in carbohydrate-deprived flies, further highlighting the specificity of this phenotype (Figure 1 – figure supplement 2H).

### *1261‐, Ir76b‐* and *Ir25a-GAL4* label common neurons required for yeast feeding

All of these lines drive expression in both olfactory (ORNs) and gustatory (GRNs) receptor neurons (Figure 1C and Figure 1 – figure supplement 2D-F). ORNs were labelled on the antennae, but not the maxillary palps (A, Figure 1C; and Figure 1 – figure supplement 2D and E), and projected to multiple non-overlapping sets of glomeruli of the antennal lobes (AL, Figure 1D and E). GRNs were labelled on the labellum (L) and in pharyngeal taste organs (LSO, VCSO), as well as on the legs (Figure 1C, Figure 1 – figure supplement 2D and E, Supplementary Table 2). *1261-GAL4* exhibited more sparse expression in GRNs than the other lines, including in pharyngeal GRNs (Supplementary Table 2). GRN projections could be seen in the subesophageal zone (SEZ, Figure 1D and E), as well as in leg and wing neuropils of the ventral nerve cord (VNC, FL, ML, HL, W, Figure 1 – figure supplement 2F).

Since all of these *GAL4* lines label chemosensory neurons, we hypothesized that all three lines label an overlapping population of sensory neurons required for yeast feeding. To test this, we generated an *Ir76b-GAL80* transgene to suppress expression of effectors in *Ir76b*-expressing neurons. Indeed, combining *Ir76b-GAL80* with *1261-GAL4* or *Ir25a-GAL4* suppressed expression in a subset of chemosensory neurons (Figure 1E and Figure 1 – figure supplement 2I). Removal of these overlapping neurons abolished the phenotype of silencing these lines, indicating that these three lines label a common population of *Ir76b*- and *Ir25a*-expressing neurons required for yeast feeding. Since *Ir25a-GAL4* exclusively labels peripheral neurons, the blockade of yeast feeding is very likely due to a sensory deficit. These lines therefore provide an entry point to dissect the sensory basis of yeast feeding.

Yeast is the main source of AAs for wild *Drosophila*, and recently a subset of *Ir76b*-expressing neurons in the legs has been proposed to respond to AAs through the Ir76b receptor^62^. We also observed a loss of preference for AA-rich food upon silencing the lines we identified: silencing *1261-GAL4*, *Ir25a-GAL4* or *Ir76b-GAL4* abolished the preference of protein-deprived flies for a diet containing AAs over one containing sucrose^26^ (Figure 1 – figure supplement 3A). However, as with other yeast metabolites (Figure 1 – figure supplement 1A), flies showed a much lower level of feeding on AAs compared to yeast (Figure 1 – figure supplement 3B). Furthermore, mutations in *Ir76b*, which abolish AA responses^62^, or in *Ir25a* had no effect on yeast feeding (Figure 1 – figure supplement 3C-E). These data strongly suggest that AAs are not the sole stimuli mediating yeast appetite.

### Taste receptor neurons within the identified lines are required for yeast feeding

Upon further inspection, we noted that all of the lines identified above show strong expression in multiple subsets of GRNs (Figure 1 and Figure 1 – figure supplement 2). We therefore hypothesized that the reduction in yeast feeding we identified would depend on silencing of taste receptor neurons common to these lines. To test this, we took advantage of the gustatory-specific enhancer of *Poxn* to generate a GRN-specific *GAL80* line (*Poxn-GAL80*). This *Poxn-GAL80* line has the unique property that, unlike the endogenous *Poxn transcript* or *Poxn-GAL4*, it is expressed broadly in the gustatory system, including taste peg GRNs. Therefore, in combination with the identified drivers, this transgene blocks the expression of effector genes in GRNs innervating taste sensilla, taste pegs, and almost all pharyngeal GRNs, while leaving expression in ORNs largely unaffected (Figure 2A, Figure 2 – figure supplement 1A and B, and Supplementary Table 2). Relieving the silencing of GRNs in any of the *GAL4* lines rescued flies’ preference for yeast, suggesting that these lines label GRNs that are necessary for yeast feeding (Figure 2A and Figure 2 – figure supplement 1A and B). To directly test the possible involvement of olfactory receptor neurons (ORNs) labelled by these lines, we performed multiple loss-of-function manipulations of ORNs. In accordance with the above results, none of the manipulations of ORNs, including *atonal* mutants which lack all *Ir*-expressing ORNs, had any effect on yeast preference (Figure 2 – figure supplement 1C-G). This suggests that blockade of ORNs labelled in these lines does not account for the yeast feeding phenotype of silencing the *GAL4* lines identified above, and indicates that this phenotype is due to a loss of gustatory input.

**Figure 2.**
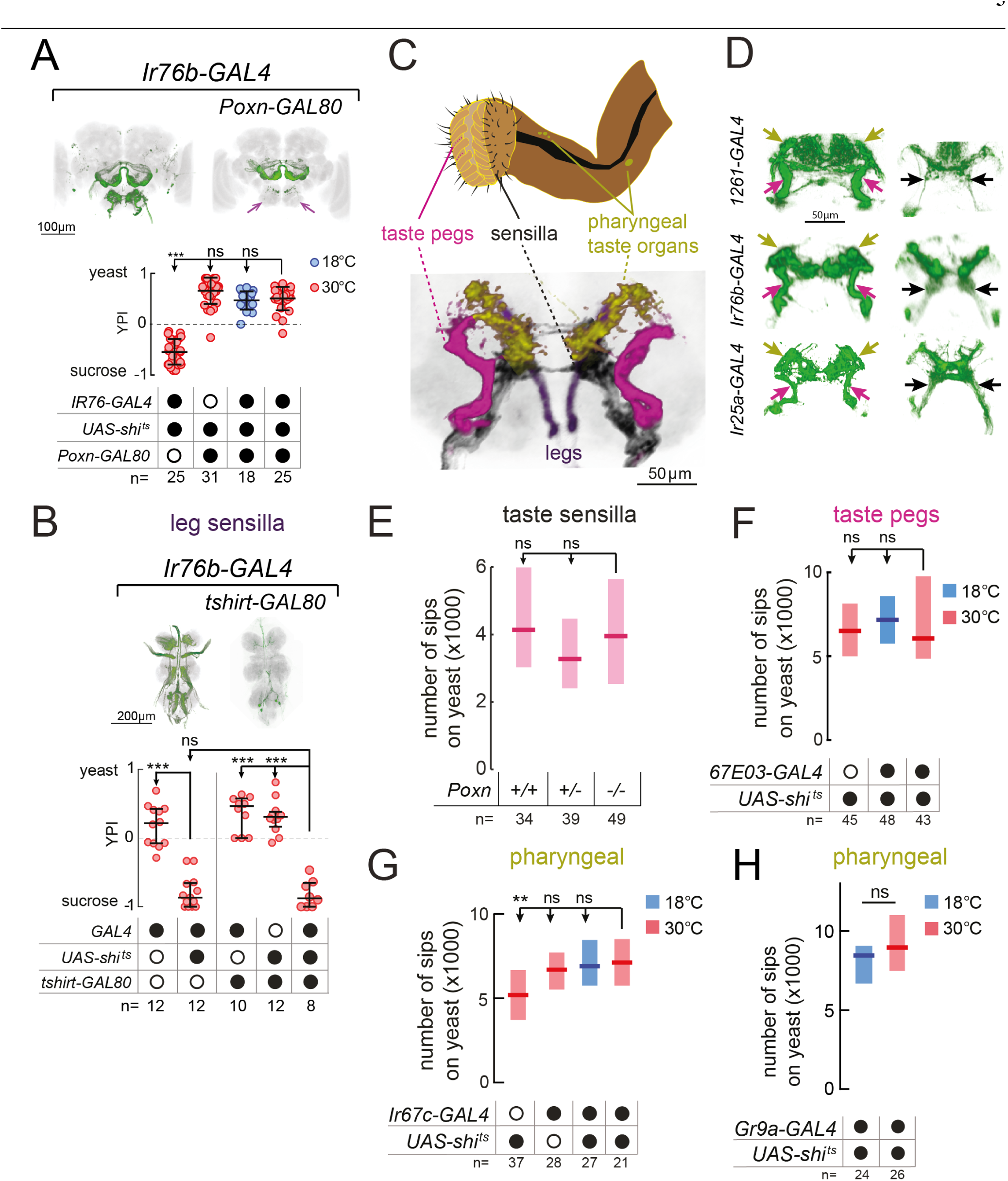
A subset of gustatory receptor neurons is required for yeast intake. (A) YPI of yeast-deprived female flies in which *Ir76b-GAL4* neurons were all silenced, or with *Poxn-GAL80*, and corresponding controls. Expression pattern of experimental flies in the brain as visualized using *UAS-CD8::GFP* in green, with nc82 synaptic staining in gray. Arrows indicate loss of GFP expression in the SEZ. (B) YPI of yeast-deprived female flies in which *Ir76b-GAL4* neurons were all silenced, or with *tshirt-GAL80*, and corresponding controls. Expression pattern of experimental flies in the VNC as visualized using *UAS-CD8::GFP* in green, with nc82 synaptic staining in gray. (C) Upper: Schematic of the proboscis, showing sensilla, taste pegs and pharyngeal taste organs. Lower: Schematic view of the SEZ of flies expressing CD8::GFP from a fragment of the Ir76b enhancer (*VT033654-GAL4*) with GRNs coloured by their peripheral innervation and gray representing the nc82 synaptic staining. Pink, taste pegs; black, sensilla; yellow, pharyngeal taste organs; purple, legs. (D) Expression of *1261*‐, *Ir76b*‐ and *Ir25a-GAL4* in the anterior (left) and posterior (right) SEZ, with arrows showing projections from pharyngeal (yellow), taste peg (pink) and sensillar (black) GRNs. (E) Number of sips from yeast by flies with 0/1/2 copies of the *Poxn*^∆*M22-B5*^ mutation. The homozygous mutant also contains a rescue construct to rescue all defects except taste sensilla (see Methods). (F-H) Number of sips from 10% yeast by yeast-deprived females in which *67E03*‐ (F), *Ir67c*‐ (G) or *Gr9a-GAL4* (H) neurons were acutely silenced, and corresponding controls. *67E03-GAL4* labels taste peg GRNs, *Ir67c-GAL4* labels a subset of GRNs in the LSO, and *Gr9a-GAL4* labels a subset of GRNs in the VCSO. In (A) and (B), circles represent yeast preference in single assays, with line representing the median and whiskers the interquartile range. In (E-H), boxes represent median with upper/lower quartiles. ***p<0.001, **p<0.01, *p<0.05, ns p≥0.05. Groups compared by Kruskal-Wallis test, followed by Dunn’s multiple comparison test.

### Proboscis gustatory receptor neurons in distinct locations act in parallel to support yeast feeding

All of the lines we identified label GRNs in both the legs and the proboscis. To test the role of leg GRNs in the observed phenotypes, we used *tshirt-GAL80* to remove tarsal and wing GRNs from the expression pattern of *Ir76b-GAL4*^68^(Figure 2B). This did not suppress the phenotype of silencing this line, indicating that the GRNs required for yeast feeding reside in the proboscis.

Within the proboscis, GRNs are present in 3 types of taste organ: in taste sensilla on the external surface of the labellum; in taste pegs on the inner surface; and in pharyngeal taste organs that contact food after ingestion41 (Figure 2C). GRNs innervating these distinct structures send axonal projections to distinct regions of the SEZ^42,69–71^. Within all of the lines we identified, we observed projections in the PMS4 region, which receives input largely from phagostimulatory sensillar GRNs (black arrows); the AMS1 region, which receives input largely from taste peg GRNs (pink arrows); and in the dorso-anterior SEZ, which receives pharyngeal GRN input (green arrows, Figure 2D).

We next aimed to separate the role of these distinct taste structures by manipulating each in turn. To remove taste sensilla function, we used a mutation in *Poxn*^72^. We found that flies lacking taste sensilla fed on yeast to the same extent as controls, indicating that sensillar GRNs alone are dispensable for yeast feeding (Figure 2E). Furthermore, silencing of neurons expressing *Poxn-GAL4*^72^, which labels GRNs innervating sensilla and the LSO, had no effect on yeast feeding, but strongly reduced feeding on sucrose, further reinforcing the evidence that the processing of these sensory modalities is separated (Figure 2 – figure supplement 2A-C).

In order to gain access to taste peg GRNs, we visually inspected a database of *GAL4* lines^73^ and identified two lines, *67E03*‐ and *57F03-GAL4*, which show expression in taste peg GRNs in addition to other central neurons^56^ (Figure 2 – figure supplement 2D). We used these lines to silence taste peg GRNs, but, as with our manipulation of sensillar GRNs, observed no effect on yeast feeding (Figure 2F and Figure 2 – figure supplement 2E). Likewise, silencing *Ir56d-GAL4*, which labels taste peg GRNs as well as a subset of sensillar GRNs^53,70^, did not affect yeast feeding (Figure 2 – figure supplement 2F). These data are in agreement with the lack of yeast feeding phenotype observed upon silencing of either *E409-GAL4* or *Gr64e-GAL4*, which have previously been characterized to label taste peg gustatory neurons^54,63^ (Figure 1 – figure supplement 1B and C). Likewise, combining *67E03-GAL4* with *Gr5a-GAL4*, which labels a subset of sensillar GRNs^42,44^, was not sufficient to decrease yeast feeding (Figure 2 – figure supplement 2G).

Finally, to manipulate pharyngeal GRNs, we conducted a series of experiments in which we blocked output from subsets of *Gr*‐ and *Ir*-expressing GRNs previously shown to innervate the different pharyngeal taste organs^70,71^ (Figure 2G and H, and Figure 2 – figure supplement 3). None of these lines showed a reduction in yeast feeding, suggesting that the tested pharyngeal GRNs are not essential to support yeast feeding.

Taken together, these data indicate that the GRNs in the proboscis common to the three identified lines play an important role in mediating yeast feeding, and support the idea that the identified GRNs innervating sensilla, taste pegs and pharyngeal organs act redundantly in mediating yeast feeding, such that if one set is compromised, the others still suffice to support yeast feeding. Given that yeast is an essential resource for *Drosophila*, a strategy which relies on distributed gustatory structures to ensure yeast intake would be highly advantageous. For this to be true, one would predict that different subsets of GRNs within the identified lines should respond to the taste of yeast.

### *Ir76b* labellar GRNs respond to yeast taste

We have shown that *Ir76b*‐ and *Ir25a*-expressing GRNs in the proboscis of the fly are required for yeast intake. To characterize the response of these neurons to yeast taste, we performed two-photon calcium imaging of *Ir76b* GRN axons in the SEZ of awake flies while stimulating the labellum with liquid stimuli (Figure 3A). To capture the response in all the projection fields of GRN axons, we imaged at multiple planes spanning the antero-posterior extent of the SEZ (Figure 3B). In agreement with the behavioral data above, we observed strong calcium responses to yeast in both the AMS1 and PMS4 regions, where axons of neurons innervating the taste pegs and taste sensilla, respectively, terminate^69^ (Figure 3C and D). These yeast responses were significantly greater than responses to water or 500 mM sucrose (Figure 3E and F), and were sharply aligned to stimulus onset and offset (Figure 3 – figure supplement 1A and B). *Ir76b*-positive taste peg and sensillar neurons therefore preferentially respond to yeast taste, supporting the view that multiple subsets of GRNs on the labellum are involved in yeast feeding.

**Figure 3.**
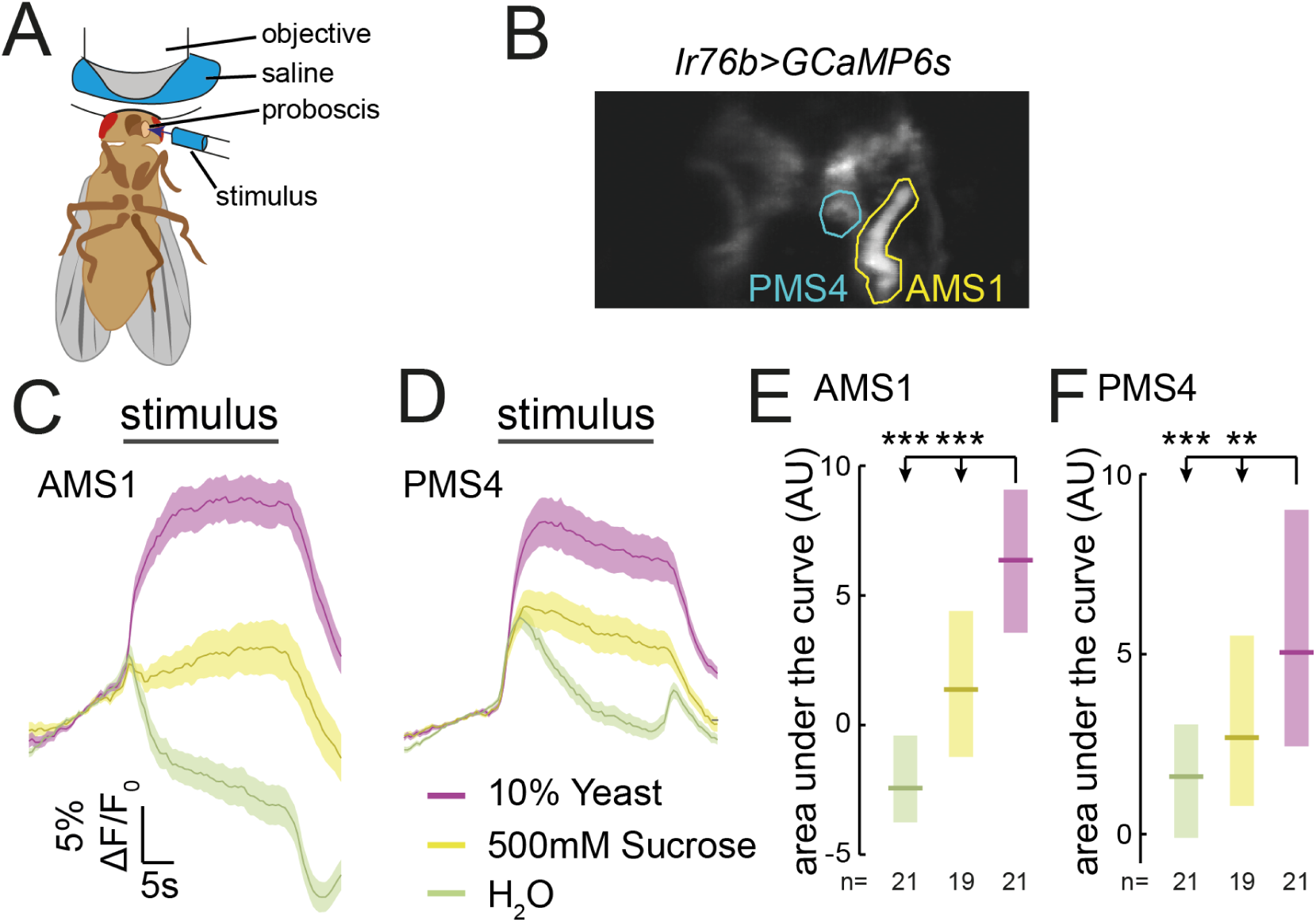
*Ir76b* GRNs respond to yeast taste. (A) Schematic of the imaging setup. The fly is head-fixed, with a window in the head to allow visual access to GRN axons in the SEZ. The fly is stimulated on the labellum with liquid tastant solutions. (B) Representative z-projection of baseline GCaMP6s fluorescence from *Ir76b-GAL4* axons in the SEZ. Highlighted ROIs indicate the AMS1 and PMS4 regions, which are largely innervated by taste peg and sensillar GRNs, respectively. (C-F) Average responses measured in ROIs shown in (B) to 10% yeast (purple), 500 mM sucrose (yellow) or water (green) from females deprived of yeast for 10 days. Average (mean +/-SEM) trace of ∆F/F_0_ from GCaMP6s signal in AMS1 (C) and PMS4 (D) upon taste stimulation. Black line indicates stimulus period. Responses quantified as area under the curve in AMS1 (E) and PMS4 (F) during stimulus presentation. Boxes represent median with upper/lower quartiles. **p<0.01, ***p<0.001. Groups compared by Kruskal-Wallis test, followed by Dunn’s multiple comparison test.

The response to yeast in AMS1 is in agreement with earlier data indicating that taste peg GRNs respond to carbonation, a by-product of yeast metabolism^63^. We confirmed that taste peg GRNs responded to carbonation; however, we found that these GRNs also responded to deactivated yeast, which does not produce carbonation, suggesting that taste peg GRNs can detect yeast independently from carbonation (Figure 3 – figure supplement 1C). Likewise, the response in PMS4 is consistent with reported activation of sensillar GRNs by glycerol^54,55^, though the response to glycerol concentrations in the range found in yeast are significantly smaller than the response to yeast^74^ (Figure 3 – figure supplement 1D). Furthermore, we observed an increase in calcium in response to putrescine in PMS4, but not in AMS1 (Figure 3 – figure supplement C and D). These data indicate that yeast metabolites activate distinct subsets of *Ir76b*-positive neurons, which is consistent with our behavioral data indicating that silencing individual subsets of yeast GRNs does not affect yeast feeding (Figure 2).

### The response of yeast GRNs is modulated by internal amino acid state

In order to direct feeding decisions towards achieving nutrient homeostasis, animals must integrate information about their current needs with sensory information from foods available in the environment. Mated female flies, when deprived from dietary yeast, respond with a homeostatic increase in yeast feeding^27^ (Figure 4A and Figure 1 – figure supplement 2A). This appetite is yeast-specific, since sucrose feeding is not increased by yeast deprivation (Figure 4C). Flies are therefore able to adjust their food intake in a nutrient-specific fashion. We speculated that internal nutrient state changes, induced by deprivation from yeast, may specifically affect the response to yeast taste at the level of sensory neurons. To test this, we imaged the responses of *Ir76b* GRNs to yeast taste in flies fed on sucrose alone for varying durations. We found that 3 days of yeast deprivation resulted in a small but nonsignificant increase in the response of both taste peg and sensillar GRNs to yeast, and that this response was significantly enhanced after 10 days of yeast deprivation (Figure 4B and Figure 4 – figure supplement 1A). The responses of *Ir76b* GRNs to the water solvent, in contrast, were not increased by yeast deprivation (Figure 4 – figure supplement 1B and C).

**Figure 4.**
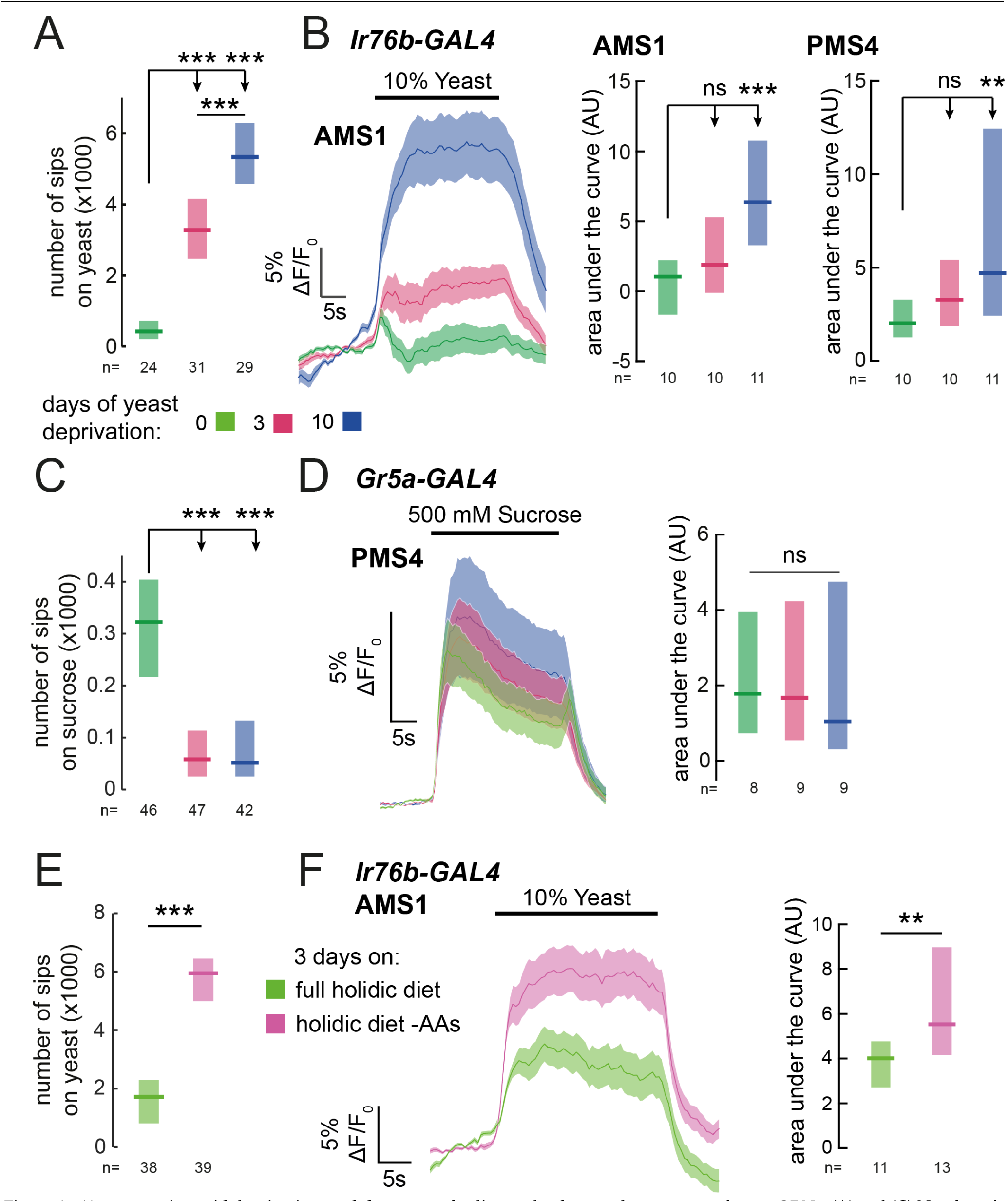
Yeast or amino acid deprivation modulates yeast feeding and enhances the response of yeast GRNs. (A) and (C) Number of sips from 10% yeast (A) and 20 mM sucrose (C) by female flies of the genotype *Ir76b-GAL4, UAS-GCaMP6s.* Flies were deprived from yeast for 0, 3 or 10 days. (B) Left: Average (mean +/-SEM) trace of ∆F/F0 from GCaMP6s signal in *Ir76b-GAL4* AMS1 region upon stimulation of the labellum with 10% yeast, from flies deprived of yeast for 0, 3 or 10 days. Quantification of responses of AMS1 (center) and PMS4 (right) regions as area under the curve during stimulus presentation. (D) Left: Average (mean +/-SEM) trace of ∆F/F0 measured from GCaMP6s in *Gr5a-GAL4* taste sensillar projections upon stimulation of the labellum with 500 mM sucrose, from flies deprived of yeast for 0, 3 or 10 days. Right: Quantification of responses as area under the curve during stimulus presentation. (E) Number of sips from 10% yeast by females fed on a holidic diet with or without amino acids for 3 days prior to assay. (F) Left: Average (mean +/-SEM) trace of ∆F/F0 from GCaMP6s signal in *Ir76b-GAL4* AMS1 region upon stimulation of the labellum with 10% yeast, from flies treated as in (E). Right: Quantification of responses as area under the curve during stimulus presentation. (A), (C) and (E) Feeding behaviour of mated female flies expressing GCaMP6s under the control of *Ir76b-GAL4* was measured on the flyPAD. Boxes represent median with upper/lower quartiles. ***p<0.001, **p<0.01, *p<0.05, ns p≥0.05. (A-D) Groups compared by Kruskal-Wallis test, followed by Dunn’s multiple comparison test. (E-F) Groups compared by Wilcoxon rank-sum test.

If this sensory modulation is part of a nutrient-specific appetite, it should be specific to GRNs that support feeding on yeast, and not other taste stimuli. Indeed, we found that the response of sugar-sensing GRNs (*Gr5a-GAL4*) to sucrose was not enhanced by yeast deprivation (Figure 4D). In fact, the response to the low concentration of sucrose used in the behavioral experiments was suppressed by this yeast deprivation regime, mirroring the effect of yeast deprivation on sucrose feeding (Figure 4C and Figure 4 – figure supplement 1D). This indicates that yeast deprivation is distinct from starvation, which has been shown to increase the sugar sensitivity of Gr5a neurons^66^. Thus, deprivation from a specific food, yeast, specifically enhances the sensitivity of primary sensory neurons detecting that nutrient source, providing a potential basis for homeostatic changes in nutrient choice.

Of all the nutrients present in yeast, AAs are the most potent modulators of reproduction and lifespan, critical life-history traits of the animal^20^. It is therefore unsurprising that internal AA state is the main nutritional factor regulating yeast appetite^26^. We hypothesized that the increase in sensory gain to yeast would be driven by the internal AA state of the fly, such that GRNs would have increased gain when the fly is low on AAs. Alternatively, exposure to dietary yeast could desensitize yeast GRNs, so that a diet devoid of yeast would increase their sensitivity independently of the flies’ nutritional state. To disentangle the effects of AA state from sensory experience, we turned to a chemically defined diet, which allowed us to manipulate specific components of the diet independently, in the absence of exposure to yeast^20,25,75^. As expected, removal of AAs from the flies’ diet elicited a specific appetite for yeast (Figure 4E). We then imaged the response of yeast-sensing GRNs in flies deprived specifically from AAs, and found that the gain of these GRNs is enhanced by AA deprivation (Figure 4F). These data indicate that the internal AA state of the fly modulates the gain of yeast-sensing GRNs, potentially allowing the fly to homeostatically compensate for the lack of AAs.

### Reproductive state acts downstream of sensory neurons to modulate yeast feeding

Animals have to constantly integrate information from multiple internal states to produce adaptive behaviors. Accordingly, yeast appetite is regulated not only by AA state, but also by the reproductive state of the fly^25,27–29^. Mating disinhibits yeast appetite through a dedicated circuit, such that mated females consume significantly more yeast than virgins^29^ (Figure 5A). Our finding that yeast feeding is mediated by specific GRNs strongly suggests that as with salt taste^29^, mating changes the behavioral response to yeast taste. We therefore hypothesized that the mating state of the animal could be integrated with nutrient state to drive yeast appetite in three distinct manners: by directly modulating the response of yeast-sensing GRNs; by modulating the sensitivity of the nervous system to protein deprivation; or by modulating downstream processing independently of AA state (Figure 5B). Having a cellular readout for the effect of these internal states on yeast taste responses allows us to distinguish these hypotheses. We imaged the response of yeast GRNs in virgin and mated females in different yeast deprivation regimes. While yeast deprivation had a strong impact on taste peg and sensillar GRN responses to yeast, there was no significant effect of mating on these responses (Figure 5C and D). This is in contrast to the synergistic interaction between deprivation and mating states seen in the flies’ feeding behavior, and suggests that reproductive state may act on downstream gustatory processing to influence flies’ feeding on yeast. In support of this hypothesis, we found that knockdown of *Sex Peptide Receptor* in *Ir76b*-expressing neurons had no effect on females’ postmating yeast appetite (Figure 5 – figure supplement 1A), reinforcing the concept that Sex Peptide acts through the SPSN-SAG pathway, and not directly on yeast-sensing neurons, to stimulate yeast appetite in anticipation of the demands of egg production^27,29,30^. Furthermore, by pooling together virgin and mated female imaging data, we now found that the response of yeast-sensing GRNs was significantly increased following just 3 days of protein deprivation (AMS1: p=0.004; PMS4: p=0.005, AUC). Taken together, these data show that the nervous system has independent mechanisms for detecting nutritional and reproductive states, and that these two states are integrated independently at different levels of the yeast feeding circuit.

**Figure 5.**
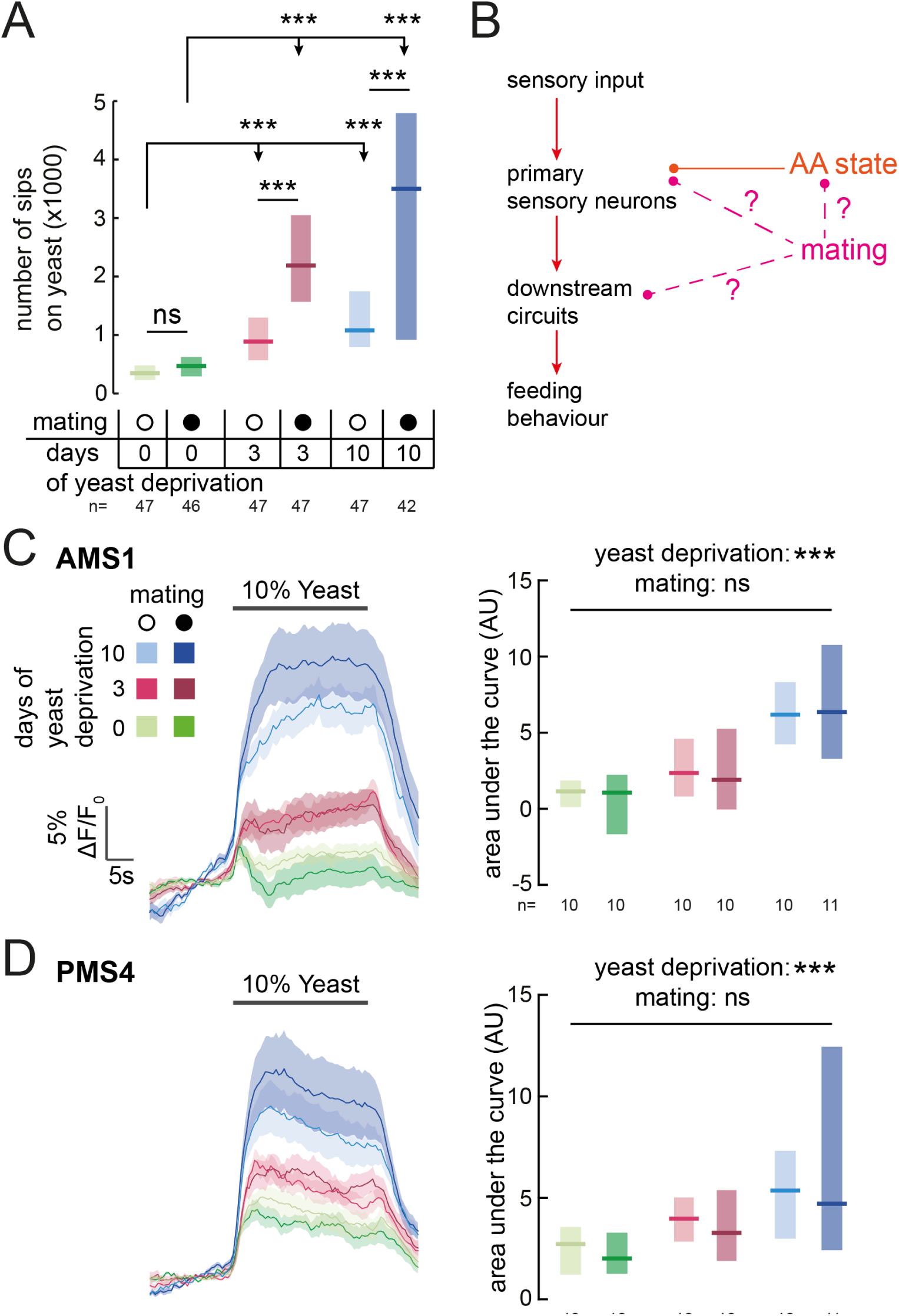
Mating state affects yeast feeding behavior but does not modulate GRN responses to yeast. (A) Number of sips from 10% yeast by female flies of the genotype *Ir76b-GAL4,UAS-GCaMP6s*. Flies were virgin or mated, and yeast-deprived for 0, 3 or 10 days. (B) Schematic of how mating could act to modulate yeast feeding behavior. Mating could regulate the sensitivity of the nervous system to AA deprivation; it could act on yeast GRNs independently from protein state; or it could act on downstream yeast taste processing circuits. (C) and (D) Calcium responses of AMS1 (C) and PMS4 (D) regions of *Ir76b-GAL4* upon stimulation of the labellum with 10% yeast, from virgin and mated females deprived of yeast for 0, 3 or 10 days. Left: Average (mean +/-SEM) trace of ∆F/F0. Right: Quantification of responses as area under the curve during stimulus presentation. Boxes represent median with upper/lower quartiles. Significance was tested using two-way ANOVA, with deprivation and mating states as the independent variables. In (A), this was followed by multiple comparisons with Bonferroni correction. ***p<0.001, ns p≥0.05.

## DISCUSSION

The utility of environmental resources to animals is dependent on their current internal states. As such, animals must adapt their decision-making depending on these states, choosing resources that fulfil current organismal requirements while minimizing the negative impact of mismatches. In the context of feeding, this means not only regulating caloric intake, but also balancing the intake of different nutrients1. In this study, we identify resource-specific modulation of primary sensory neurons as a potential mechanism underlying such state-dependent tuning of value-based decisions. Specifically, we show that the amino acid (AA) state of the fly modulates the gain of gustatory sensory neurons that mediate the ingestion of yeast, flies’ main source of dietary protein, such that the response of these neurons to yeast is increased when the fly is lacking AAs. The response of sensory neurons detecting sugars, in contrast, is not increased by protein deprivation, indicating that this modulation is resource-specific. Rather, the response of sugar-sensing neurons is increased by complete starvation^66^. Thus, the gain of these two classes of attractive taste-coding neurons is separately regulated to drive nutrient-specific appetites.

Yeast appetite is driven by the lack of dietary essential AAs^20,25,26^, and AA state modulates GRN responses to yeast. However, the mechanism through which AA state regulates GRN gain is not known and will be an important avenue for future research. AA state could act at two levels in yeast GRNs: at presynaptic terminals within the SEZ, or at the peripheral level, to adapt GRN responses to the internal state. Recent studies in *Drosophila* have indicated that complete starvation state acts through distinct neuromodulatory mechanisms on the presynaptic terminals of sensory neurons in the brain to increase attraction to food odors^76^ and sugars^65,66^ and to decrease sensitivity to bitter deterrents^77,78^. Intriguingly, acute blockade of synaptic release from these neuromodulatory systems did not suppress yeast appetite induced by yeast deprivation (Supplementary Table 1), suggesting that a distinct mechanism may regulate sensory gain according to AA state. Alternatively, this modulation could occur at the level of peripheral sensory responses, as seen for AA responses in the fish olfactory and locust gustatory systems^79,80^. In locusts, this peripheral modulation occurs through desensitization by AAs in the hemolymph. Regardless of the mechanism, our results demonstrate nutrient-specific modulation of the gain of select sensory neuron responses as an elegant way to increase the salience of the specific resources that are important to maintain homeostasis in the current state, and thus to optimize lifespan and reproduction. AA state is likely to act at multiple levels in the nervous system, in addition to its effect on primary sensory neurons, to drive a robust yeast appetite ‐similarly to the multi-level integration seen in other systems^81^. Recently, two central systems have been proposed to modulate protein appetite in *Drosophila*: the dopaminergic WED-FB-LAL circuit, which promotes yeast intake in protein-deprived flies^9^, and the fat body-derived hormone FIT, which acts in the brain to suppress intake of protein-rich food in the sated state^8^. Identifying whether these systems modulate sensory processing, or affect other aspects of the complex behavioral changes that guide protein homeostasis^25^, will be important for a mechanistic understanding of the circuit basis of protein appetite. The dynamics of value-based decisions are shaped by multiple ongoing internal states, which must interact in the nervous system. How different states interact at the circuit level to shape behavior is poorly understood. Feeding on yeast is known to be synergistically modulated by both AA state and reproductive state^25,27–29^. Here, we demonstrate that these two states are separately detected by the nervous system and influence taste processing at different levels of the sensory circuit. While AA state strongly modulates the gain of neurons mediating yeast feeding, reproductive state has no effect on these responses. Although virgin and mated females show very different behavioral responses to protein deprivation, our results suggest that they are subject to the same lack of AAs, and that their sensory neurons are modulated in a similar way by protein deprivation. Mating is therefore likely to gate downstream gustatory processing to enhance yeast appetite in anticipation of the nutritional demands of egg production^30^. This result stands in contrast to the previously-described modulation of *Ir76b*+ GRN responses to polyamines in the AMS1 region that occurs transiently following mating^82^. This modulation is dependent on SPR expression in GRNs; however, in the context of yeast feeding, we show that SPR knockdown in *Ir76b*+ neurons has no effect (Figure 5 – figure supplement 1). Rather, mating state is detected by Sex Peptide sensory neurons in the reproductive tract, and conveyed to the brain by SAG neurons to modulate yeast and salt appetite, with the postmating yeast appetite also requiring the action of octopamine^25,29,83^. How the activity of SAG neurons modulates downstream gustatory processing, however, remains to be discovered.

Yeast is a critical component of the diet of *Drosophila* melanogaster, providing most dietary proteins in addition to many other non-caloric nutrient requirements^21,24^. Here, we identify a population of gustatory receptor neurons that is necessary for yeast feeding and responds to yeast taste. These *Ir76b*‐ and *Ir25a*-expressing neurons are distributed across the proboscis. The strong yeast feeding phenotype, the anatomical overlap of the lines we identified, as well as the strong and selective response of these neurons to yeast, and the fact that they are regulated by yeast deprivation, indicates that the identified neurons play a key role in yeast intake. Manipulation of specific subsets of these neurons did not affect yeast feeding, suggesting that they act in a redundant manner to support yeast feeding. This suggests that the detection of this important resource relies on a distributed set of gustatory neurons in the proboscis of the fly.

Yeast is a complex, multimodal resource, and thus it is likely that multiple stimuli present in yeast activate *Drosophila* chemosensory neurons^37,54,63^. At long ranges, flies are attracted by yeast volatiles detected through the olfactory receptor neurons^58–60^. Olfaction is also important for efficient recognition of yeast as a food source^25^. However, we show that input from the gustatory system is ultimately critical to elicit feeding on yeast. Multiple yeast fermentation products, including carbonation, activate the gustatory neurons identified in this study. However, these stimuli on their own do not induce a feeding rate that approximates the phagostimulatory power of yeast. Rather, our results suggest that multiple yeast stimuli must coincide to produce a yeast percept, and that yeast GRNs are likely to be specialized to respond to a variety of chemicals normally found in this microorganism. This would allow flies to ensure the reliable detection of this essential food source, while permitting selective regulation of feeding on yeast depending on internal state. The emerging picture is that multiple yeast metabolites are detected by different gustatory neurons on the proboscis which act redundantly to mediate yeast intake, thus forming a proxy for the perception and ingestion of protein-, and therefore AA-, rich food.

Intriguingly, some *Ir76b*+ GRNs in the legs have been shown to respond to AAs^62^. However, we show here that tarsal GRNs are dispensable for yeast feeding, and do not contribute to the *Ir76b-GAL4* silencing phenotype. Furthermore, Ir20a, the putative receptor that conveys AA responsiveness, is not expressed in labellar GRNs^70^, and flies lacking *Ir76b*, which is required for these AA responses, feed on yeast to the same extent as controls. It is therefore possible that AA taste does not significantly contribute to the detection of yeast but that flies are also able to detect this important nutrient independently to ensure their uptake from non-yeast sources. Furthermore, it is interesting to note that *Ir76b* GRNs have been shown to mediate the taste of substances which are important to support reproduction^29,37,51,62^. *Ir76b* GRNs could therefore be specialized in mediating the intake of foods that are relevant for reproduction. Overall, our data demonstrate that deprivation from a particular nutrient – AAs – specifically increases the gain of gustatory neurons detecting the ethological food substrate – yeast – that provides the animal with this nutrient. In principle, such nutrient-specific sensory gain modulation could represent a general mechanism through which internal states could change the salience of a resource by directing behaviors towards stimuli that are most relevant in a specific state.

## Acknowledgements

We thank Richard Benton, Gerry Rubin, Kristin Scott, Leslie Vosshall, Ilona Grunwald-Kadow, Craig Montell, Scott Waddell, Ann-Shyn Chiang, Amita Sehgal, Aaron DiAntonio, Elizabeth Gavis, Michael Pankratz, Irene Miguel-Aliaga, Bader Al-Anzi, Ping Shen, Werner Boll, Barry Dickson, Sofia Lavista-Llanos and Gero Miesenböck for providing fly strains. We also thank Ulrike Heberlein, Wes Grueber, the NP consortium, Douglas Armstrong, and numerous other researchers that have contributed to our collection of GAL4 stocks. Further lines obtained from the Bloomington Drosophila Stock Center (NIH P40OD018537), and the VDRC (Vienna, Austria) were used in this study. The enhancer trap GAL4 silencing screen was performed at the IMP in the laboratory of Barry J. Dickson. We thank Barry J. Dickson, Martin Häsemeyer, Nilay Yapici, Christoph Treiber, Lorenz Pammer, Carolina Doran, and Ana Paula Elias for assistance during the screen and post-hoc analysis. We thank Richard Benton, Juan Antonio Sánchez-Alcañiz and Kristin Scott for training and assistance in establishing our two-photon calcium imaging prep. We thank Eugenia Chiappe, Michael Orger, Dennis Goldschmidt, Daniel Münch, and members of the Behavior and Metabolism laboratory for helpful discussions and comments on the manuscript. This project was supported by the Portuguese Foundation for Science and Technology (FCT) grant PTDC/BIA-BCM/118684/2010, and postdoctoral fellowship SFRH/BPD/79325/2011 to PMI; the Human Frontier Science Program Project Grant RGP0022/2012; the BIAL Foundation Grant (283/14); and the Marie Curie FP7 Programme FLiACT (ITN) grant. Research at the Centre for the Unknown is supported by the Champalimaud Foundation.

### AUTHOR CONTRIBUTIONS

Conceived and developed the project: CR, SJW, KS; Performed the screens: CR, KS, CB; Performed two-photon imaging experiments: SJW; Performed intersectional genetic analysis of circuits, neuroanatomy, and behavioral experiments: KS, SJW, CB, CR; Performed data analysis and interpretation: KS, SJW, PMI, CR; Developed and built hardware and developed software: SJW, PMI; Wrote the manuscript: SJW, CR, KS.

## METHODS

### Fly Husbandry

All data are from yeast-deprived mated female flies unless otherwise stated. Flies were reared at 18 or 25°C, 70% relative humidity on a 12 h light-dark cycle. Experimental and control flies were reared at standard density and were matched for age and husbandry conditions. The fly medium (yeast-based medium [YBM]) contained, per litre, 80 g cane molasses, 22 g sugar beet syrup, 8 g agar, 80 g corn flour, 10 g soya flour, 18 g yeast extract, 8 ml propionic acid, and 12 ml nipagin (15% in ethanol). For experiments that did not involve thermogenetic silencing (including calcium imaging), flies were kept at 25°C. Yeast deprivation was induced by feeding flies for 3 (or 10) days on a tissue soaked with 6.5 ml of 100 mM sucrose. For nutrient-specific deprivation, flies were reared on yeast-based food, transferred to fresh YBM for one day and then kept for 3 days on holidic medium with or without amino acids. This holidic medium was prepared as detailed in^20,25^, using the 50S200NYaa composition to approximate the amino acid ratio found in yeast. For experiments involving thermogenetic silencing, flies were reared and kept at 18°C. Yeast deprivation was induced by keeping flies for 7 days on a tissue soaked with 100 mM sucrose. This longer deprivation time was chosen to compensate for the lower metabolic rate at colder temperatures.

### *Drosophila* stocks and genetics

Neuronal silencing was achieved using the 20xUAS-shibirets transgene inserted in the VK00005 or attP5 landing site (gift of Gerry Rubin, HHMI Janelia Research Campus). Genetic backgrounds of the control animals were matched as closely as possible to the experimental animals. For silencing experiments, the VK00005 or attP5 landing site background was crossed to the GAL4 line as a control. The *Poxn* mutant fly experiments contained the following genetic manipulations (gift of Markus Noll and Werner Boll): homozygous control: *w*^1118^; heterozygous control: *w*^1118^; *Poxn*^∆M22-B5^/+; homozygous mutant: *w*^1118^; *Poxn*^∆M22-B5^ homozygote with *∆SfoBs105*/*∆SfoBs127* to rescue all defects other than taste sensilla. The atonal mutant fly experiments relied on the following genotypes (gift of Ilona Kadow): Control flies (*ato*^+^/^+^): *ey*^*FLP*^; *FRT82B/FRT82B, CL*; Flies with atonal mutant antennae (*a*to^-/-^): *ey*^*FLP*^; *FRT82B ato*[1]/*FRT82B, CL*. The full genotypes of the lines used in the manuscript are listed in Supplementary Table 3.

The *Poxn-GAL80* vector was a gift of Duda Kvitsiani and Barry Dickson. Briefly, the 14kb *Poxn* enhancer fragment from the *Poxn-GAL4-14* vector^72^ was cloned into a custom *GAL80*-containing vector, which was injected into *w*^1118^ embryos.

To generate *Ir76b-GAL80*, we amplified the enhancer fragment of *Ir76b* using the following primers: 5’-CCCAGTCTAATGTATGTAATTGCC, 5’-CGATACGAGTGCCTACTGTAC, and cloned into the pBPGAL80Uw-6 vector (AddGene). This vector was separately inserted into the attp40 and attP2 sites by PhiC31 integrase-mediated recombination. Injections were performed by BestGene.

### Neuronal silencing experiments

Flies carrying the temperature-sensitive allele of *shibire* under UAS control were crossed with a collection of different *GAL4* lines in order to acutely silence distinct neuronal subpopulations in experimental flies. Flies were reared at 18°C. 7-10 days after eclosion, female flies were sorted into fresh YBM and Canton-S males were added to ensure mating. Two days later, flies were transferred again to YBM; on the following day, they were transferred to 100 mM sucrose solution for 7 days to induce a yeast deprivation state. Silencing was induced by pre-incubating flies at 30°C for two hours and performing the two-color food choice or the flyPAD assays at this temperature. Control flies were always kept at 18°C, including during the assay.

### Two-color food choice assays

Two-color food choice assays were performed as previously described^27^. Groups of 16 female and 5 male flies were briefly anesthetized using light CO_2_ exposure and introduced into tight-fit lid petri dishes (Falcon, #351006). For the yeast choice assays, the flies were given the choice between nine 10 μl sucrose spots mixed with red colorant (20 mM sucrose [Sigma-Aldrich, #84097]; 7.5 mg/ml agarose [Invitrogen, #16500]; 5 mg/ml Erythrosin B [Sigma-Aldrich, #198269]; 10% PBS) and nine 10 μl spots of yeast mixed with blue colorant (10% yeast [SAF-instant, Lesaffre]; 7.5 mg/ml agarose; 0.25 mg/ml Indigo carmine [Sigma-Aldrich, #131164]; 10% PBS) for 2 hours at 18, 25 or 30°C, 70% RH, depending on the experimental condition. Flies were then frozen and females scored by visual inspection as having eaten either sucrose (red abdomen), yeast (blue abdomen), or both (red and blue or purple abdomen) media. The yeast preference index (YPI) for the whole female population in the assay was calculated as follows: (n_blue yeast_ - n_red sucrose_)/(n_red sucrose_ + n_blue yeast_ + n_both_). For amino acid preference experiments, yeast was replaced with a holidic diet (200NHunt) lacking sucrose, and sucrose was replaced with a holidic diet (50 mM sucrose) lacking amino acids, as described in^26^. The holidic diet was prepared as detailed in^20,25^ according to the Hunt amino acid ratio. Both solutions were in 1% agarose. Values for n shown in the figures indicate the number of food choice assays performed.

### flyPAD assays

flyPAD assays were performed based on a protocol previously described^67^. One well of the flyPAD was filled with 20 mM sucrose, and the other with 10% yeast, each in 1% agarose. In Figure 1 – figure supplement 1A and 3B, only one well of the flyPAD was filled with the stimulus solution in 1% agarose. Deactivated yeast was generated using an autoclave; carbonation was produced by mixing 0.2 ml of 1 M NaHCO_3_ with 0.8 ml of 1 M NaH_2_PO_4_ immediately before use^63^; glycerol (Sigma-Aldrich, #G6279) was used at 10 mM (approximately the concentration in live yeast^74^) and putrescine (Sigma-Aldrich, #51799) at 5 mM. Flies were individually transferred to flyPAD arenas by mouth aspiration and allowed to feed for one hour at the indicated temperature, 70% RH; except for Figure 1 – figure supplement 1A and Figure 1 – figure supplement 3B, where flies fed for 30 minutes. flyPAD data were acquired using the Bonsai framework^84^, and analyzed in MATLAB using custom-written software, as described in^67^. Values for n shown in the figures indicate the number of flies tested.

### Immunohistochemistry, image acquisition and 3D rendering

Males from each *GAL4* line were crossed to females homozygous for the *UAS-CD8::GFP* reporter line and 3-10 day-old adult females heterozygous for the *GAL4* driver and UAS reporter were dissected. Samples were dissected in 4°C PBS and were then transferred to Formaldehyde solution (4% paraformaldehyde in PBS + 10% Triton-X) and incubated for 20-30 minutes at RT. Samples were washed 3 times in PBST (0.3% TritonX in PBS) and then blocked in Normal Goat Serum 10% in PBST for 2-4 hours at RT. Samples were then incubated in primary antibody solutions (Rabbit anti-GFP [Torrey Pines Biolabs] at 1:2000 and Mouse NC82 [Developmental Studies Hybridoma Bank] at 1:10 in 5% Normal Goat Serum in PBST). Primary antibody incubations were performed overnight at 4°C with rocking. They were then washed in PBST 2-3 times for 10-15 minutes at RT and again washed overnight at 4°C. The secondary antibodies were applied (Anti-mouse A594 [Invitrogen] at 1:500 and Anti-rabbit A488 [Invitrogen] at 1:500 in 5% Normal Goat Serum in PBST) and brains were then incubated for 3 days at 4°C. They were again washed in PBST 2-3 times for 10-15 minutes at RT, and washed overnight at 4°C. Before mounting the samples were washed for 5-10 minutes in PBS. Samples were mounted in Vectashield (Vector Laboratories). Images were captured on a Zeiss LSM 710 using 10x or 20x objectives. Images of the periphery did not include immunostainings. Heads and legs of female flies were clipped off and placed in a drop of Oil 10S (VWR chemicals) between a slide and a cover slip. Images were captured on a Zeiss LSM 710 using a 10x objective. Note that during the mounting procedure of the fly head, the labellum opens, exposing the inner surface of the labellum and the taste pegs. 3D reconstructions of the nervous system and the periphery were generated using FluoRender^85,86^.

### Calcium imaging

The preparation for calcium imaging was adapted from that described in^87^. Each fly was lightly anaesthetized using CO_2_, and fixed into a custom-made chamber using UV-curing glue (Bondic). The proboscis was fixed by the rostrum in an extended position, and UV-curing glue was used to seal around the head capsule within the imaging window, which was then immersed in AHL saline (103 mM NaCl, 3 mM KCl, 5 mM TES, 10 mM trehalose dihydrate, 10 mM glucose, 2 mM sucrose, 26 mM NaHCO_3_, 2 mM CaCl2 dihydrate, 4 mM MgCl2 hexahydrate, 1 mM NaH2PO4, pH7.3) bubbled with 95%O2/5%CO2. A 30G needle (BD Microlance 3) was used to cut the cuticle along the edges of the eyes, just above the rostrum and just below the ocelli, and this piece of cuticle along with the antennae was removed using forceps. Air sacs and fatty tissue surrounding the SEZ were removed, but the esophagus was left intact. The stage was then transferred to a mount under a two-photon resonant-scanning microscope (Scientifica) equipped with a 20x NA 1.0 water immersion objective (Olympus). During imaging, the brain was constantly perfused with AHL saline bubbled with 95%O2/5%CO_2_.

A 920nm laser (Coherent) was used to excite GCaMP6s through a resonant scanner, and emitted fluorescence was recorded using a photomultiplier tube. The objective was controlled by a piezo-electric z-focus, allowing serial volumetric scans of the SEZ. The SEZ was imaged at 31 z-positions, with the upper and lower limits defined in order to encompass the entire SEZ (~60-80 μm), at a frame rate of 61.88 Hz, with a 235x117 μm (512x256 pixel) frame. Imaging data were acquired using SciScan (Scientifica). A frontal view of the fly, illuminated by scattered light from the laser, was simultaneously acquired through a PointGrey Flea3 camera using Bonsai. Each trial consisted of 4650 frames (150 volumes); the gustatory stimulus was applied to the labellum at approximately volume #51 and removed at volume #100 using a pulled glass micropipette filled with the stimulus solution mounted on a micromanipulator (Sensapex). Each stimulus was applied 2-3 times per fly, and the mean response per fly used for further analysis. For comparing responses across dietary conditions, imaging sessions using flies of each condition were interleaved. Stimuli were dissolved or suspended in water as follows: 10% w/v SAF-INSTANT yeast; 10% yeast deactivated using an autoclave; 1% dry ice; 200 mM NaHCO_3_ in 500 mM NaH_2_PO_4_ (mixed immediately before use); 10 and 100 mM glycerol; 10 and 100 mM putrescine; 20 and 500 mM sucrose.

### Calcium imaging analysis

All analysis of imaging data was performed using custom scripts in MATLAB (Mathworks). To facilitate analysis of imaging data, we took the average intensity projection of each serial volume stack (31 frames). A 5x5 pixel median filter was applied to each such projection, and each projection was registered to the first projection collected from that fly to correct for movement, using a discrete Fourier transform-based subpixel rigid registration algorithm^88^. For each fly, regions of interest (ROIs) were manually defined based on the maximum intensity projection of the first trial for this fly. Two-photon and camera images were aligned using the onset of scattered laser illumination in the camera image. Each trial was then aligned to stimulus onset based on manual annotation of stimulus application time. For each trial and ROI, the mean fluorescence intensity within the ROI at each time point was used as F, and F_0_ calculated as the median of these values from 22 to 2 frames before the annotated stimulus onset. ∆F/F_0_ was calculated as (F-F_0_)/F_0_ for each time point. To calculate area under the curve, we took the sum of the ∆F/F_0_ values from 0 to 44 frames after stimulus onset for each fly’s average trace, except for Figure 3 – figure supplement 1C and D, for which frames 0 to 19 were used due to the shorter stimulation time. For each fly, we treated each brain hemisphere as a separate data point; values for n shown in the figures indicate the number of flies imaged per condition.

### Statistics

Results of two-color food choice assays, flyPAD and calcium imaging experiments were compared using the Kruskal-Wallis test, followed by Dunn’s multiple comparison test. For tests comparing only two groups, the Wilcoxon rank-sum test was used. To analyze the effects of mating and protein deprivation, groups were compared by two-way ANOVA. All tests were two-tailed.

## REFERENCES

1. Simpson, S. J. & Raubenheimer, D. The Nature of Nutrition: A Unifying Framework from Animal Adaptation to Human Obesity. (Princeton University Press, 2012).

2. Hughes, B. O. & Wood-Gush, D. G. A specific appetite for calcium in domestic chickens. Anim. Behav. 19, 490–499 (1971).

3. Deutsch, J. A., Moore, B. O. & Heinrichs, S. C. Unlearned specific appetite for protein. Physiol. Behav. 46, 619–624 (1989).

4. Beauchamp, G. K., Bertino, M., Burke, D. & Engelman, K. Experimental sodium depletion and salt taste in normal human volunteers. Am. J. Clin. Nutr. 51, 881–889 (1990).

5. Itskov, P. M. & Ribeiro, C. The dilemmas of the gourmet fly: The molecular and neuronal mechanism of feeding and nutrient decision making in *Drosophila*. Front. Decis. Neurosci. 7:, 12 (2013).

6. Jarvie, B. C. & Palmiter, R. D. HSD2 neurons in the hindbrain drive sodium appetite. Nat. Neurosci. 20, 167–169 (2017).

7. Matsuda, T. et al. Distinct neural mechanisms for the control of thirst and salt appetite in the subfornical organ. Nat. Neurosci. 20, 230–241 (2017).

8. Sun, J. et al. *Drosophila* FIT is a protein-specific satiety hormone essential for feeding control. Nat. Commun. 8,ncomms14161 (2017).

9. Liu, Q. et al. Branch-specific plasticity of a bifunctional dopamine circuit encodes protein hunger. Science 356, 534–539 (2017).

10. Dethier, V. G. Behavioral aspects of protein ingestion by the blowfly Phormia regina Meigen. Biol. Bull. 121, 456–470 (1961).

11. Simpson, S. J. & Abisgold, J. D. Compensation by locusts for changes in dietary nutrients: behavioural mechanisms. Physiol. Entomol. 10, 443–452 (1985).

12. Judson, C. L. Feeding and oviposition behavior in the mosquito *Aedes aegypti* (l.). i. Preliminary studies of physiological control mechanisms. Biol. Bull. 133, 369–377 (1967).

13. Brown, M. R. et al. Endogenous regulation of mosquito host-seeking behavior by a neuropeptide. J. Insect Physiol. 40, 399–406 (1994).

14. Griffioen-Roose, S. et al. Protein status elicits compensatory changes in food intake and food preferences. Am. J. Clin. Nutr. 95, 32–38 (2012).

15. Gosby, A. K. et al. Testing Protein Leverage in Lean Humans: A Randomised Controlled Experimental Study. PLOS ONE 6, e25929 (2011).

16. Levine, M. E. et al. Low Protein Intake Is Associated with a Major Reduction in IGF-1, Cancer, and Overall Mortality in the 65 and Younger but Not Older Population. Cell Metab. 19, 407–417 (2014).

17. Solon-Biet, S. M. et al. The Ratio of Macronutrients, Not Caloric Intake, Dictates Cardiometabolic Health, Aging, and Longevity in Ad Libitum-Fed Mice. Cell Metab. 19, 418–430 (2014).

18. Lee, K. P. et al. Lifespan and reproduction in *Drosophila*: New insights from nutritional geometry. Proc. Natl. Acad. Sci. 105, 2498–2503 (2008).

19. Grandison, R. C., Piper, M. D. W. & Partridge, L. Amino-acid imbalance explains extension of lifespan by dietary restriction in *Drosophila*. Nature 462, 1061–1064 (2009).

20. Piper, M. D. W. et al. A holidic medium for *Drosophila* melanogaster. Nat. Methods (2013). doi:10.1038/nmeth.2731

21. Baumberger, J. P. A nutritional study of insects, with special reference to microörganisms and their substrata. J. Exp. Zool. 28, 1–81 (1919).

22. Phaff, H. J., Miller, M. W., Recca, J. A., Shifrine, M. & Mrak, E. M. Yeasts Found in the Alimentary Canal of *Drosophila*. Ecology 37, 533–538 (1956).

23. De Camargo, R. & Phaff, H. J. Yeasts Occurring in *Drosophila* Flies and in Fermenting Tomato Fruits in Northern California. J. Food Sci. 22, 367–372 (1957).

24. Starmer, W. T. & Lachance, M.-A. Chapter 6 - Yeast Ecology. in The Yeasts (Fifth Edition) (eds. Kurtzman, C. P., Fell, J. W. & Boekhout, T.) 65–83 (Elsevier, 2011). doi:10.1016/B978-0-444-52149-1.00006-9

25. Corrales-Carvajal, V. M., Faisal, A. A. & Ribeiro, C. Internal states drive nutrient homeostasis by modulating exploration-exploitation trade-off. eLife 5, e19920 (2016).

26. Leitão-Gonçalves, R. et al. Commensal bacteria and essential amino acids control food choice behavior and reproduction. PLOS Biol. 15, e2000862 (2017).

27. Ribeiro, C. & Dickson, B. J. Sex Peptide Receptor and Neuronal TOR/S6K Signaling Modulate Nutrient Balancing in *Drosophila*. Curr. Biol. 20, 1000–1005 (2010).

28. Vargas, M. A., Luo, N., Yamaguchi, A. & Kapahi, P. A Role for S6 Kinase and Serotonin in Postmating Dietary Switch and Balance of Nutrients in D. melanogaster. Curr. Biol. 20, 1006–1011 (2010).

30. Walker, S. J., Goldschmidt, D. & Ribeiro, C. Craving for the future: the brain as a nutritional prediction system. Curr. Opin. Insect Sci. (2017). doi:10.1016/j.cois.2017.07.013

29. Walker, S. J., Corrales-Carvajal, V. M. & Ribeiro, C. Postmating Circuitry Modulates Salt Taste Processing to Increase Reproductive Output in *Drosophila*. Curr. Biol. CB 25, 2621–2630 (2015).

31. Dethier, V. The hungry fly: A physiological study of the behavior associated with feeding. (Harvard U Press, 1976).

32. Yarmolinsky, D. A., Zuker, C. S. & Ryba, N. J. P. Common Sense about Taste: From Mammals to Insects. Cell 139, 234–244 (2009).

33. Yang, C., Belawat, P., Hafen, E., Jan, L. Y. & Jan, Y.-N. *Drosophila* Egg-Laying Site Selection as a System to Study Simple Decision-Making Processes. Science 319, 1679–1683 (2008).

34. Joseph, R. M., Devineni, A. V., King, I. F. G. & Heberlein, U. Oviposition preference for and positional avoidance of acetic acid provide a model for competing behavioral drives in *Drosophila*. Proc. Natl. Acad. Sci. 106, 11352–11357 (2009).

35. Joseph, R. M. & Heberlein, U. Tissue-specific Activation of a Single Gustatory Receptor Produces Opposing Behavioral Responses in *Drosophila*. Genetics (2012). doi:10.1534/genetics.112.142455

36. Yang, C.-H., He, R. & Stern, U. Behavioral and Circuit Basis of Sucrose Rejection by *Drosophila* Females in a Simple Decision-Making Task. J. Neurosci. 35, 1396–1410 (2015).

37. Hussain, A. et al. Ionotropic Chemosensory Receptors Mediate the Taste and Smell of Polyamines. PLOS Biol. 14,e1002454 (2016).

38. Mann, K., Gordon, M. D. & Scott, K. A Pair of Interneurons Influences the Choice between Feeding and Locomotion in *Drosophila*. Neuron 79, 754–765 (2013).

39. Thoma, V. et al. Functional dissociation in sweet taste receptor neurons between and within taste organs of Drosophila. Nat. Commun. 7, 10678 (2016).

40. Auer, T. O. & Benton, R. Sexual circuitry in *Drosophila*. Curr. Opin. Neurobiol. 38, 18–26 (2016).

41. Stocker, R. F. The organization of the chemosensory system in *Drosophila melanogaster*: a review. Cell Tissue Res. 275, 3–26 (1994).

42. Wang, Z., Singhvi, A., Kong, P. & Scott, K. Taste Representations in the *Drosophila* Brain. Cell 117, 981–991 (2004).

43. Fujishiro, N., Kijima, H. & Morita, H. Impulse frequency and action potential amplitude in labellar chemosensory neurones of *Drosophila melanogaster*. J. Insect Physiol. 30, 317–325 (1984).

44. Marella, S. et al. Imaging Taste Responses in the Fly Brain Reveals a Functional Map of Taste Category and Behavior. Neuron 49, 285–295 (2006).

45. Weiss, L. A., Dahanukar, A., Kwon, J. Y., Banerjee, D. & Carlson, J. R. The Molecular and Cellular Basis of Bitter Taste in *Drosophila*. Neuron 69, 258–272 (2011).

46. Soldano, A. et al. Gustatory-mediated avoidance of bacterial lipopolysaccharides via TRPA1 activation in *Drosophila*. eLife 5, e13133 (2016).

47. Du, E. J. et al. Nucleophile sensitivity of *Drosophila* TRPA1 underlies light-induced feeding deterrence. eLife 5, e18425 (2016).

48. Miyamoto, T., Chen, Y., Slone, J. & Amrein, H. Identification of a *Drosophila* Glucose Receptor Using Ca^2+^ Imaging of Single Chemosensory Neurons. PLoS ONE 8, e56304 (2013).

49. Inoshita, T. & Tanimura, T. Cellular identification of water gustatory receptor neurons and their central projection pattern in Drosophila. Proc. Natl. Acad. Sci. U. S. A. 103, 1094–1099 (2006).

50. Cameron, P., Hiroi, M., Ngai, J. & Scott, K. The molecular basis for water taste in *Drosophila*. Nature 465, 91–95 (2010).

51. Zhang, Y. V., Ni, J. & Montell, C. The Molecular Basis for Attractive Salt-Taste Coding in *Drosophila*. Science 340, 1334–1338 (2013).

52. Masek, P. & Keene, A. C. Drosophila Fatty Acid Taste Signals through the PLC Pathway in Sugar-Sensing Neurons. PLoS Genet 9, e1003710 (2013).

53. Tauber, J. M. et al. A subset of sweet-sensing neurons identified by IR56d are necessary and sufficient for fatty acid taste. bioRxiv 174623 (2017). doi:10.1101/174623

54. Wisotsky, Z., Medina, A., Freeman, E. & Dahanukar, A. Evolutionary differences in food preference rely on Gr64e, a receptor for glycerol. Nat. Neurosci. 14, 1534–1541 (2011).

55. LeDue, E. E., Chen, Y.-C., Jung, A. Y., Dahanukar, A. & Gordon, M. D. Pharyngeal sense organs drive robust sugar consumption in *Drosophila*. Nat. Commun. 6,(2015).

56. Yapici, N., Cohn, R., Schusterreiter, C., Ruta, V. & Vosshall, L. B. A Taste Circuit that Regulates Ingestion by Integrating Food and Hunger Signals. Cell 165, 715–729 (2016).

57. Murata, S., Brockmann, A. & Tanimura, T. Pharyngeal stimulation with sugar triggers local searching behavior in *Drosophila*. J. Exp. Biol. jeb.161646 (2017). doi:10.1242/jeb.161646

58. Becher, P. G. et al. Yeast, not fruit volatiles mediate *Drosophila melanogaster* attraction, oviposition and development. Funct. Ecol. (2012).

59. Christiaens, J. F. et al. The fungal aroma gene ATF1 promotes dispersal of yeast cells through insect vectors. Cell Rep. 9, 425–432 (2014).

60. Dweck, H. K. M., Ebrahim, S. A. M., Farhan, A., Hansson, B. S. & Stensmyr, M. C. Olfactory Proxy Detection of Dietary Antioxidants in *Drosophila*. Curr. Biol. 25, 455–466 (2015).

61. Croset, V., Schleyer, M., Arguello, J. R., Gerber, B. & Benton, R. A molecular and neuronal basis for amino acid sensing in the Drosophila larva. Sci. Rep. 6, 34871 (2016).

62. Ganguly, A. et al. A Molecular and Cellular Context-Dependent Role for Ir76b in Detection of Amino Acid Taste. Cell Rep. 18, 737–750 (2017).

63. Fischler, W., Kong, P., Marella, S. & Scott, K. The detection of carbonation by the *Drosophila* gustatory system. Nature 448, 1054–1057 (2007).

64. Jiao, Y., Moon, S. J., Wang, X., Ren, Q. & Montell, C. Gr64f Is Required in Combination with Other Gustatory Receptors for Sugar Detection in *Drosophila*. Curr. Biol. 18, 1797–1801 (2008).

65. Marella, S., Mann, K. & Scott, K. Dopaminergic Modulation of Sucrose Acceptance Behavior in *Drosophila*. Neuron 73, 941–950 (2012).

66. Inagaki, H. K. et al. Visualizing neuromodulation in vivo: TANGO-mapping of dopamine signaling reveals appetite control of sugar sensing. Cell 148, 583–595 (2012).

67. Itskov, P. M. et al. Automated monitoring and quantitative analysis of feeding behaviour in *Drosophila*. Nat. Commun. 5,(2014).

68. Clyne, J. D. & Miesenböck, G. Sex-Specific Control and Tuning of the Pattern Generator for Courtship Song in *Drosophila*. Cell 133, 354–363 (2008).

69. Miyazaki, T. & Ito, K. Neural architecture of the primary gustatory center of *Drosophila melanogaster* visualized with *GAL4* and *LexA* enhancer-trap systems. J. Comp. Neurol. 518, 4147–4181 (2010).

70. Koh, T.-W. et al. The *Drosophila* IR20a clade of ionotropic receptors are candidate taste and pheromone receptors. Neuron 83, 850–865 (2014).

71. Kwon, J. Y., Dahanukar, A., Weiss, L. A. & Carlson, J. R. A map of taste neuron projections in the *Drosophila* CNS. J. Biosci. 39, 565–574 (2014).

72. Boll, W. & Noll, M. The Drosophila *Pox neuro* gene: control of male courtship behavior and fertility as revealed by a complete dissection of all enhancers. Dev. Camb. Engl. 129, 5667–5681 (2002).

73. Jenett, A. et al. A GAL4-Driver Line Resource for *Drosophila* Neurobiology. Cell Rep. (2012). doi:10.1016/j.celrep.2012.09.011

74. Scanes, K. T. (Orange F. S. U., Hohmann, S. & Prior, B. A. Glycerol production by the yeast *Saccharomyces cerevisiae* and its relevance to wine: A review. South Afr. J. Enol. Vitic. South Afr. (1998).

75. Piper, M. D. W. et al. Matching Dietary Amino Acid Balance to the *In Silico*-Translated Exome Optimizes Growth and Reproduction without Cost to Lifespan. Cell Metab. 25, 610–621 (2017).

76. Root, C. M., Ko, K. I., Jafari, A. & Wang, J. W. Presynaptic Facilitation by Neuropeptide Signaling Mediates Odor-Driven Food Search. Cell 145, 133–144 (2011).

77. Inagaki, H. K., Panse, K. M. & Anderson, D. J. Independent, Reciprocal Neuromodulatory Control of Sweet and Bitter Taste Sensitivity during Starvation in *Drosophila*. Neuron 84, 806–820 (2014).

78. LeDue, E. E. et al. Starvation-Induced Depotentiation of Bitter Taste in *Drosophila*. Curr. Biol. 26, 2854–2861 (2016).

79. Nikonov, A. A., Butler, J. M., Field, K. E., Caprio, J. & Maruska, K. P. Reproductive and metabolic state differences in olfactory responses to amino acids in a mouth brooding African cichlid fish. J. Exp. Biol. 220, 2980–2992 (2017).

80. Simpson, S. J. & Simpson, C. L. Mechanisms Controlling Modulation by Haemolymph Amino Acids of Gustatory Responsiveness in the Locust. J. Exp. Biol. 168, 269–287 (1992).

81. Ohyama, T. et al. A multilevel multimodal circuit enhances action selection in *Drosophila*. Nature 520, 633–639 (2015).

82. Hussain, A., Üçpunar, H. K., Zhang, M., Loschek, L. F. & Kadow, I. C. G. Neuropeptides Modulate Female Chemosensory Processing upon Mating in *Drosophila*. PLOS Biol. 14, e1002455 (2016).

83. Rezával, C., Nojima, T., Neville, M. C., Lin, A. C. & Goodwin, S. F. Sexually Dimorphic Octopaminergic Neurons Modulate Female Postmating Behaviors in *Drosophila*. Curr. Biol. 24, 725–730 (2014).

84. Lopes, G. et al. Bonsai: an event-based framework for processing and controlling data streams. Front. Neuroinformatics 9,(2015).

85. Wan, Y., Otsuna, H., Chien, C.-B. & Hansen, C. An Interactive Visualization Tool for Multi-channel Confocal Microscopy Data in Neurobiology Research. IEEE Trans. Vis. Comput. Graph. 15, 1489–1496 (2009).

86. Wan, Y., Otsuna, H., Chien, C.-B. & Hansen, C. FluoRender: An Application of 2D Image Space Methods for 3D and 4D Confocal Microscopy Data Visualization in Neurobiology Research. IEEE Pac. Vis. Symp. Proc. IEEE Pac. Vis. Symp. 201–208 (2012).

87. Flood, T. F. et al. A single pair of interneurons commands the *Drosophila* feeding motor program. Nature 499, 83–87 (2013).

88. Guizar-Sicairos, M., Thurman, S. T. & Fienup, J. R. Efficient subpixel image registration algorithms. Opt. Lett. 33, 156–158 (2008).

